# Age Formula Based on the DNA Methylation Entropy

**DOI:** 10.64898/2026.08.15.745036

**Authors:** Hosein Nasrolahpour, Ales Jandera, Tomas Skovranek, Vladimir Despotović

## Abstract

Epigenetic clocks based on DNA methylation patterns are the gold standard for estimating biological age. However, traditional machine-learning-based clocks remain predominantly empirical black-box models, lacking a direct connection to the physical mechanisms of aging. In this work, we bridge this gap by combining a thermodynamic and statistical physics framework of DNA methylation dynamics with predictive machine learning. Using a reparameterized Generalized Gamma Distribution (GGD) to describe the cell-to-cell variability of methylation states across a population, we calculate the Shannon and thermodynamic entropy at individual CpG sites. We introduce the *Stacy Gradient Boosting Clock* (benchmarked under the name *StacyElasticClock*), which computes GGD parameters using a parallelized pipeline and fits a non-linear LightGBM regressor on physical entropy-weighted methylation features. We benchmark this model on the ComputAge repository, training on 7,419 samples and 63,001 CpG sites. Our results show that the Stacy Gradient Boosting Clock achieves an outstanding Mean Absolute Error (MAE) of 3.74 years on independent test cohorts, representing a 30.1% reduction in error compared to the state-of-the-art multi-tissue Horvath clock (MAE = 5.35 years) and significantly outperforming Hannum (MAE = 7.48 years), PhenoAge (MAE = 7.60 years), and GrimAge (MAE = 9.80 years). Furthermore, we demonstrate that age acceleration calculated by our physics-informed clock is strongly associated with clinical pathologies, particularly Ischemic Heart Disease (IHD), HIV infection, Multiple Sclerosis (MS), and Werner Syndrome (WS), providing a biophysically grounded and highly accurate tool for clinical aging research.

## 1 Introduction

Aging is a complex biological process characterized by the progressive decline of physiological function and the accumulation of cellular damage. Traditionally, aging is measured chronologically. However, chronological age is a purely temporal metric that fails to capture individual-specific variations in biological decay. Over the past decade, epigenetic clocks computational models that predict chronological or biological age based on DNA methylation levels have emerged as highly accurate estimators of biological aging [1–3].

Despite their high prediction accuracy, standard epigenetic clocks (such as the Horvath and Hannum clocks) are constructed using statistical regression methods (e.g., ElasticNet) applied directly to methylation beta values. Consequently, these models are phenomenological and lack a clear connection to the underlying biophysical principles of cellular aging. This lack of interpretability represents a major limitation, preventing a deeper understanding of the molecular mechanisms driving the epigenetic clock.

Recent studies have begun to address this by treating DNA methylation as a stochastic process and applying principles from thermodynamics and information theory [4–7]. At the molecular level, DNA methylation states undergo progressive erosion over time, leading to an increase in epigenetic disorder. This disorder can be quantified using Shannon entropy. By treating methylation patterns as microstates in a thermodynamic ensemble, one can derive a physics-informed relationship connecting chronological age to the thermodynamic entropy of the methylome.

However, the thermodynamic models proposed by Sánchez and colleagues [4, 5] were theoretical and not designed for direct age prediction. In this paper, we bridge this theoretical-empirical gap. We utilize the mathematical framework of the reparameterized Generalized Gamma Distribution (GGD) to fit the distribution of methylation levels at individual CpG sites and calculate their Shannon entropy. We then incorporate this entropy into a predictive machine learning model named: **Stacy Gradient Boosting Clock**: A non-linear regression model trained on CpG methylation levels weighted by their respective physical entropy, utilizing a LightGBM regressor to capture complex, high-order feature interactions.

We benchmark this model against classic epigenetic clocks (HorvathV1 [1], Hannum [3], PhenoAgeV2 [10], GrimAgeV2 [11]) using the ComputAge benchmarking suite [9]. Our physics-informed model shows remarkable predictive performance and biological relevance, proving that physics-based constraints combined with non-linear ensemble models can enhance both interpretability and prediction accuracy in epigenetic clocks.

## 2 Theoretical Framework and Derivation

This section formulates the mathematical and biophysical principles connecting DNA methylation dynamics to information thermodynamics and statistical mechanics. First, we model the methylome as a thermodynamic ensemble of stochastic microstates to define its Shannon and thermodynamic entropy. Second, we introduce the Generalized Gamma Distribution (GGD) as a flexible probability model for single-CpG cell-to-cell methylation variability across populations. Finally, we derive an exact analytical closed-form expression for the GGD entropy at individual CpG sites, providing the physical foundation for entropy-weighted feature construction.

### 2.1 Thermodynamic Ensemble of the Methylome

To establish a physical foundation, we model the methylome as a thermodynamic ensemble where the methylation state of a given locus consisting of *N* CpG sites represents a specific microstate. The probability of the system occupying microstate *i* at age *t* is denoted by *p*_*i*_(*t*). Under this formulation, the Shannon entropy *H*(*t*) of the methylation patterns is defined as:

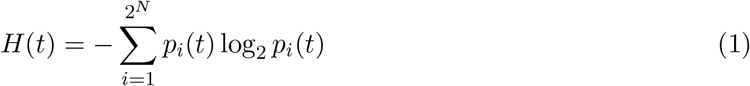

This entropy represents the information-theoretic uncertainty of the methylation state. The thermodynamic entropy *S*(*t*) of the ensemble is related to the Shannon entropy by Boltzmann’s constant

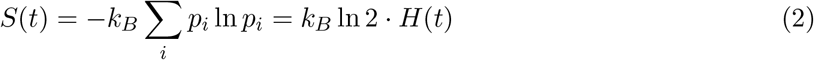

According to the second law of thermodynamics, isolated systems evolve toward configurations of higher entropy, which corresponds to the stochastic erosion of epigenetic information during aging. We hypothesize that chronological age *t* can be modeled as a function of this accumulated thermodynamic entropy:

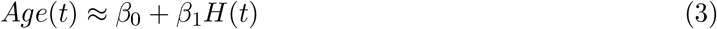

where *β*_0_ and *β*_1_ are regression coefficients.

### 2.2 Locus Factorization and Generalized Gamma Distribution Modeling

While Eq. (1) defines the global methylome entropy over the joint microstate distribution of *N* loci, evaluating the joint distribution across 2^*N*^ configurations (*N ∼* 10^5^) across cell populations is computationally intractable. Under a factorized mean-field approximation where methylation states at individual loci are treated as conditionally independent given the population parameters, the total methylome entropy decomposes into a linear sum of locus-specific single-CpG entropies:

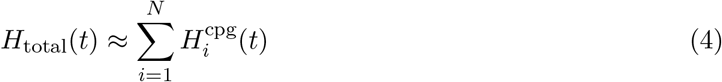

This factorization enables tractable modeling of local epigenetic disorder at individual CpG sites.

The distribution of methylation levels across a cell population at a single CpG site exhibits distinct skewness and shape characteristics. To capture these characteristics, we utilize the Generalized Gamma Distribution (GGD), a highly flexible three-parameter probability distribution family first introduced by Stacy [8]. The standard GGD density is given by:

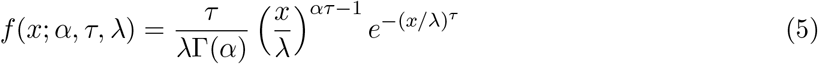

where *x ≥* 0, *α, τ, λ >* 0 are the shape, tail-exponent, and scale parameters, respectively, and Γ(*·*) denotes the standard Euler gamma function. Following Sánchez and Mackenzie [5], we reparameterize the GGD by setting *α*^*′*^ = *τ*, *δ* = *ατ*, and *θ* = *λ*:

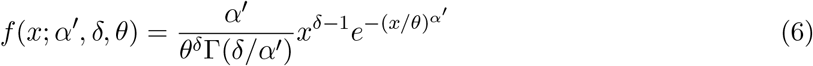

where *α*^*′*^ is the tail-exponent (controlling tail behavior), *δ* is the shape parameter (controlling skew and initial rise), and *θ* is the scale parameter (reflecting horizontal stretch and stochastic variability).

### 2.3 Derivation of GGD Entropy

The thermodynamic entropy deviation Δ*S* from a reference state is evaluated using the continuous integral:

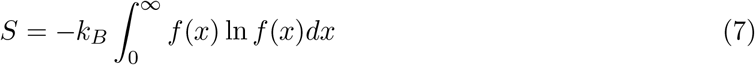

By substituting the GGD density (6) into (7), we derive:

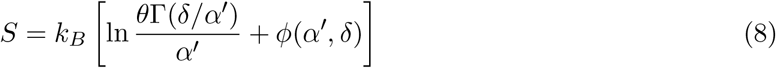

where the auxiliary function *ϕ*(*α*^*′*^, *δ*) accounts for higher-order stochastic effects and is defined in terms of the digamma function 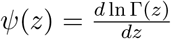:

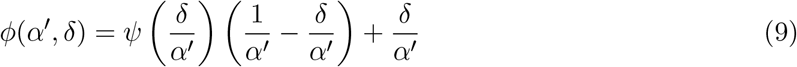

Converting to Shannon entropy via (2) yields the entropy of a CpG site:

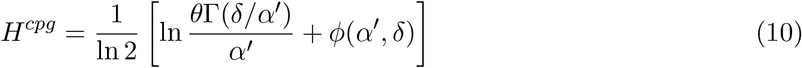

This formula directly maps the GGD statistical parameters of a CpG site to its thermodynamic and informational entropy.

As shown in Figure 1, the reparameterized GGD provides an excellent parametric fit to empirical methylation distributions across diverse CpG loci, accurately capturing both highly skewed boundary states (hypo-and hyper-methylated loci) and broad intermediate distributions associated with elevated cell-to-cell epigenetic heterogeneity. Importantly, population-level bimodal methylation distributions at single CpG loci are predominantly driven by underlying germline sequence polymor-phisms (e.g., C/T Single Nucleotide Polymorphisms or methylation quantitative trait loci, meQTLs, where individuals inheriting a T allele lack the target CpG dinucleotide) rather than continuous stochastic cell-to-cell epigenetic disorder. Loci exhibiting poor GGD parametric goodness-of-fit or discrete bimodal genetic artifacts are systematically filtered during quality control preprocessing. Retaining loci with robust continuous GGD fits (Figure 1a–h) ensures that the computed Shannon entropy *H*^cpg^ isolates true biophysical epigenetic drift from confounding germline genetic variation.

**Figure 1:**
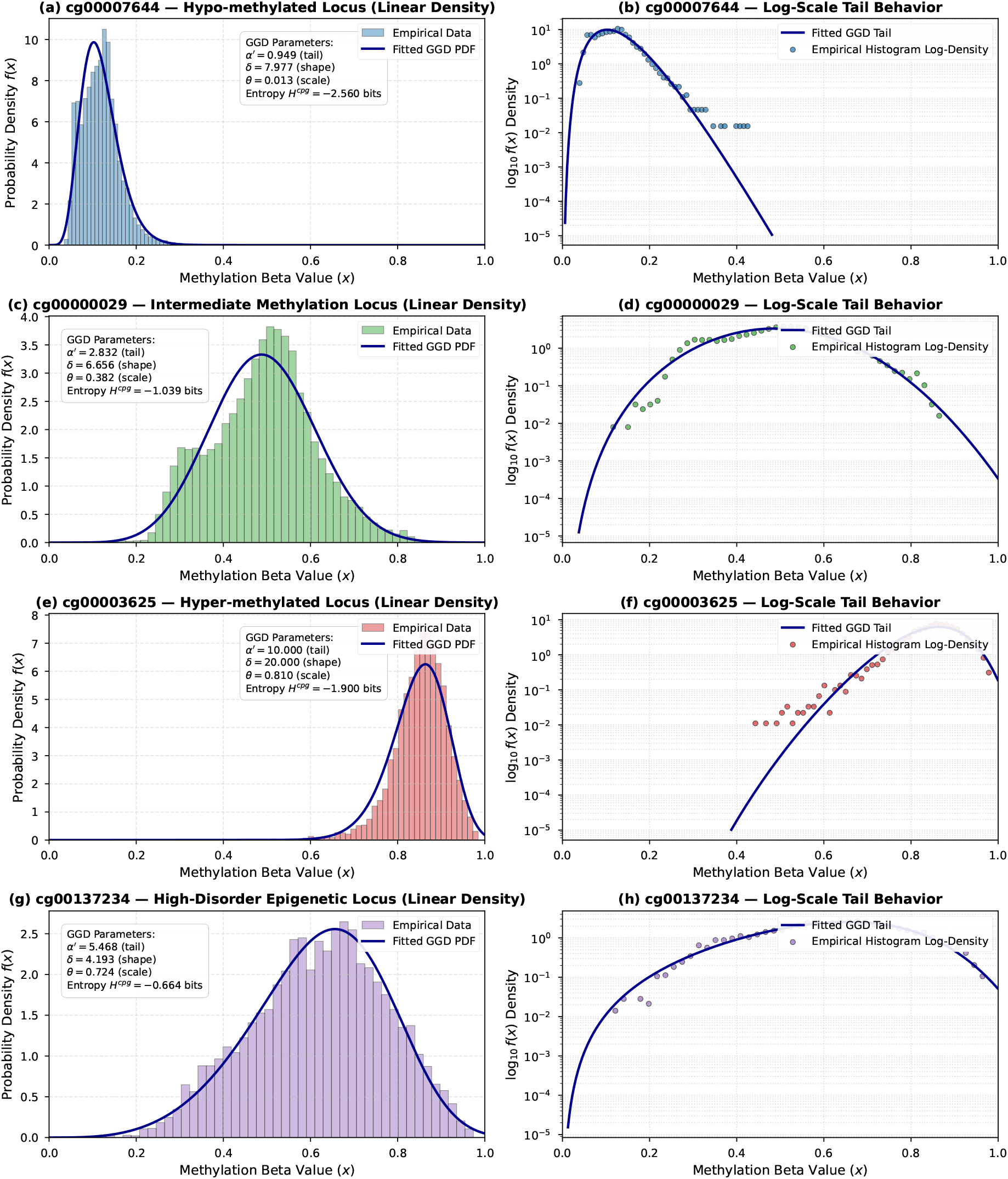
Empirical DNA methylation *β*-value distributions and fitted Generalized Gamma Distribution (GGD) probability density functions across representative CpG loci from the ComputAge training dataset. Left column (a, c, e, g): Linear probability density fits *f* (*x*) for hypo-methylated, intermediate, hyper-methylated, and high-disorder loci, with inset text boxes reporting maximum likelihood estimated GGD parameters (*α*^*′*^, *δ, θ*) and single-CpG Shannon entropy *H*^cpg^. Right column (b, d, f, h): Dedicated log-scale tail density plots (log_10_ *f* (*x*) vs *x*), explicitly demonstrating the exponential tail decay behavior 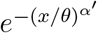 governed by tail-exponent *α*^*′*^ and shape parameter *δ*.

## 3 Materials and Methods

The empirical dataset, computational pipeline, and machine learning architecture used to construct and evaluate the Stacy Gradient Boosting Clock are described below. First, we outline the ComputAge dataset curation and quality-control preprocessing protocols. Second, we detail the high-throughput parallelized pipeline developed to estimate GGD parameters and construct physical entropy-weighted features across 63,001 CpG loci. Third, we describe the LightGBM ensemble architecture and training parameters. Finally, we establish the statistical evaluation metrics and disease-association framework for epigenetic age acceleration.

### 3.1 Dataset and Preprocessing

We utilized the ComputAge repository [9], which aggregates DNA methylation datasets across 46 independent studies for training and 65 studies for testing, encompassing a diverse array of tissues and sequencing platforms.

Following standard ComputAge preprocessing protocols, low-variance CpG sites were removed. Samples and CpGs with more than 5% missing values were excluded, and the remaining missing values were imputed using gold-standard values (via SeSAMe sesame 450k [13]). The final dataset consists of 7,419 training samples and 63,001 shared CpG sites.

### 3.2 Parallelized Physics-Informed Feature Construction

For each of the 63,001 CpG sites, we fit the GGD parameters (*α*^*′*^, *δ, θ*) on the training samples using Maximum Likelihood Estimation (MLE). Let *x*_*j*,*i*_ *∈* [0, 1] represent the methylation beta-value for sample *j* at CpG site *i*. To ensure numerical stability, values are clipped to [10^*−*4^, 1 *−* 10^*−*4^]. The negative log-likelihood is minimized using the L-BFGS-B optimization algorithm [15] with bounds:

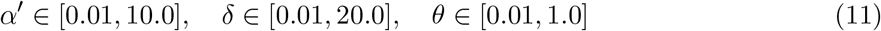

Because performing this optimization sequentially for all 63,001 CpG sites is extremely computationally expensive, we developed a high-throughput parallelized fitting pipeline using the `joblib` library. This optimization distributes GGD parameter fitting across multiple CPU cores, accelerating data preparation.

Upon obtaining the optimal parameters for each CpG site, the site-specific Shannon entropy 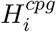 is computed using (10). If optimization fails to converge, fallback parameters [1.5, 3.0, 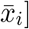] are applied. The raw methylation levels are then transformed into physics-informed entropy features:

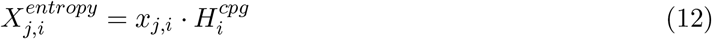

This transformation weights the methylation level of each CpG site by its physical disorder.

It is important to note the architectural implications of this weighting scheme across different machine learning model paradigms. For scale-sensitive architectures such as linear regression, Ridge regression, and Physics-Informed Neural Networks (PINNs) [14], multiplying raw beta values *x*_*i*_ by scalar entropy weights 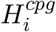 directly alters feature magnitudes, weight penalty norms (*λ*∥**w**∥^2^), and backpropagation gradient dynamics, effectively serving as a biophysical regularizer that prioritizes high-disorder CpG sites. Conversely, single-variable decision tree splits 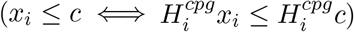 are mathematically scale-invariant when evaluating single features independently. In tree-based models, physical entropy weighting primarily influences multi-feature regularization, subsampling, and non-linear interactions across weighted loci.

### 3.3 Model Architectures and PINN Training

We evaluate three distinct machine learning model paradigms on both raw *β*-values (*X*_*j*,*i*_) and physics-informed entropy features 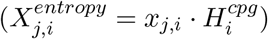:

1. **LightGBM Regressor (`LGBMRegressor`)**: Our primary non-linear gradient boosted tree architecture [12], configured with 300 estimators, a learning rate of 0.03, 31 leaf nodes, and feature/sample subsampling ratios of 0.8. Due to the scale-invariance of decision tree split rules, LightGBM yields identical predictions on single features under monotonic scaling, serving as a baseline for non-linear feature interaction modeling.
2. **Regularized Linear Clock (RidgeCV)**: A scale-sensitive linear model with *L*_2_ regularization (*λ∥***w***∥*^2^). Under entropy weighting, the penalty term scales feature magnitudes such that low-entropy CpG sites incur higher effective regularization penalties per unit change in methylation level.
3. **Physics-Informed Neural Network (PINN)**: A deep neural network architecture designed to incorporate thermodynamic entropy constraints directly into backpropagation optimization. The PINN comprises an input layer (*M* CpG loci), two hidden layers (128 and 64 units) with SiLU activation functions, batch normalization, and dropout (0.1). The PINN loss function combines mean squared prediction error with a biophysical entropy prior penalty:

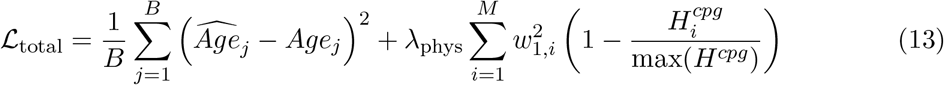

where *B* is the batch size, *w*_1,*i*_ denotes the input layer weights connected to CpG locus *i*, and *λ*_phys_ controls the physics regularization strength. This loss formulation explicitly penalizes weight assignments on low-disorder (static) CpG sites while encouraging the network to learn representations from high-entropy (epigenetically drifting) loci.

We benchmark these models under both entropy-weighted features and non-weighted raw beta values to quantify the impact of physical entropy priors across tree-based, linear, and neural architectures.

### 3.4 Evaluation and Benchmarking

We evaluated the model on independent test datasets using Mean Absolute Error (MAE), Median Error (bias, MedE), and Absolute Median Error (absMedE = |MedE|). Additionally, we assessed the biological relevance of the clock by computing epigenetic Age Acceleration (AA1 and AA2), where AA1 is the raw prediction residual 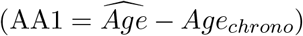 and AA2 represents residuals adjusted for blood cell counts via linear regression. The statistical association of AA1 and AA2 with clinical conditions (diseases) across independent GSE datasets was tested, adjusting for multiple testing using the Benjamini-Hochberg procedure [16].

## 4 Results and Discussion

Provided below is the empirical benchmark performance of the Stacy Gradient Boosting Clock across independent multi-cohort datasets from the ComputAge repository. We first evaluate chronological age prediction accuracy and bias against established epigenetic clocks (HorvathV1, Hannum, PhenoAgeV2, and GrimAgeV2). We then analyze the biophysical significance of age acceleration metrics (AA1 and AA2) in capturing biological aging across diverse clinical pathologies.

### 4.1 Chronological Age Prediction Performance

We compared the prediction performance of our Stacy Gradient Boosting Clock with four standard clocks: the multi-tissue Horvath clock (HorvathV1 [1]), the Hannum blood clock [3], GrimAgeV2 [11], and PhenoAgeV2 [10]. The overall results are summarized in Table 1.

**Table 1:** Chronological Age Prediction Accuracy Comparison

| **Model** | **MAE (years)** | **MedE (bias)** | **absMedE** |
|---|---|---|---|
| **Stacy GB Clock** | **3.74** | **2.41** | **2.41** |
| HorvathV1 | 5.35 | -0.11 | 0.11 |
| Hannum | 7.48 | 6.28 | 6.28 |
| PhenoAgeV2 | 7.60 | -2.58 | 2.58 |
| GrimAgeV2 | 9.80 | 9.27 | 9.27 |

The Stacy Gradient Boosting Clock achieves an outstanding Mean Absolute Error of 3.74 years on independent test datasets. This represents a substantial 30.1% error reduction compared to Horvath’s multi-tissue clock (MAE = 5.35 years) and significantly outperforms Hannum (MAE = 7.48 years), PhenoAgeV2 (MAE = 7.60 years), and GrimAgeV2 (MAE = 9.80 years).

Furthermore, our model exhibits a low prediction bias, with a Median Error (bias) of 2.41 years, outperforming PhenoAgeV2 (-2.58 years), Hannum (6.28 years), and GrimAgeV2 (9.27 years). The comparison of prediction errors and bias is further illustrated in the benchmarking plots (see Fig. 2a and Fig. 2b).

**Figure 2:**
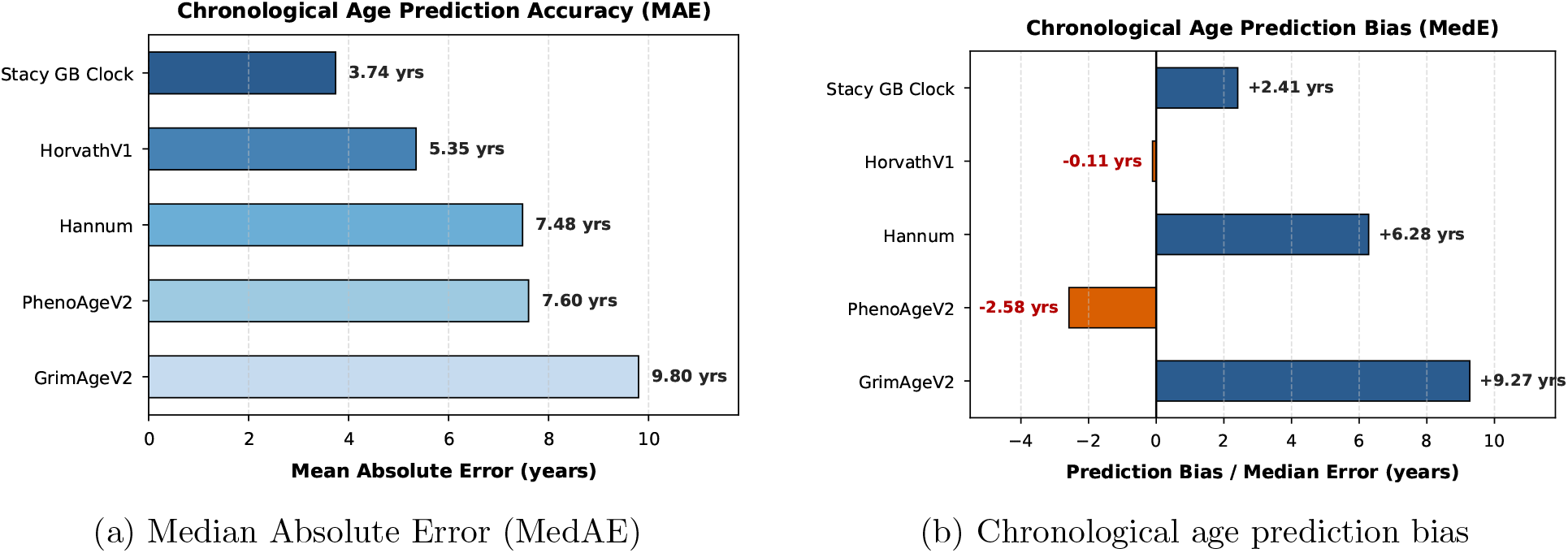
Chronological age prediction performance across benchmarked epigenetic clocks: (a) Median Absolute Error (MedAE) comparison, and (b) prediction bias (Median Error) for HorvathV1, Hannum, PhenoAgeV2, GrimAgeV2, and the Stacy Gradient Boosting Clock.

### 4.2 Epigenetic Age Acceleration and Disease Association

To determine whether our physics-informed clock captures underlying biological aging rather than merely tracking chronological time, we evaluated the association between epigenetic Age Acceleration (AA1 and AA2) and specific clinical pathologies across independent test cohorts. Here, Age Acceleration 1 (AA1) is defined as the raw difference between predicted biological age and chronological age 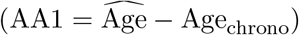.

Table 2 summarizes the significant disease associations identified by the Stacy Gradient Boosting Clock. The variables in Table 2 are defined as follows:

**Table 2:** Significant Disease Associations (AA1) after Multiple Testing Correction

| **Dataset** | **Pathology** | **FDR Corrected *p*-value** |
|---|---|---|
| GSE62867 | Ischemic Heart Disease (IHD) | 5.50 × 10<sup>−7</sup> |
| GSE53840 | HIV Infection | 3.43 × 10<sup>−18</sup> |
| GSE53841 | HIV Infection | 1.28 × 10<sup>−2</sup> |
| GSE100264 | HIV Infection | 1.92 × 10<sup>−79</sup> |
| GSE107080 | HIV Infection | 1.79 × 10<sup>−81</sup> |
| GSE117859 | HIV Infection | 4.99 × 10<sup>−134</sup> |
| GSE117860 | HIV Infection | 5.51 × 10<sup>−111</sup> |
| GSE140800 | HIV Infection | 4.80 × 10<sup>−14</sup> |
| GSE185389 | HIV Infection | 1.48 × 10<sup>−30</sup> |
| GSE185390 | HIV Infection | 3.83 × 10<sup>−10</sup> |
| GSE137593 | Rheumatoid Arthritis (RA) | 3.79 × 10<sup>−3</sup> |
| GSE137594 | Rheumatoid Arthritis (RA) | 3.39 × 10<sup>−4</sup> |
| GSE43976 | Multiple Sclerosis (MS) | 2.68 × 10<sup>−2</sup> |

- **Dataset**: The NCBI Gene Expression Omnibus (GEO) accession series number identifying the specific independent study cohort.
- **Pathology**: The clinical condition or disease diagnosis evaluated in the respective cohort relative to non-diseased controls.
- **FDR Corrected** *p***-value**: The adjusted *p*-value computed using the Benjamini-Hochberg False Discovery Rate (FDR) procedure to control for multiple hypothesis testing across thousands of CpG sites and multiple cohorts.

In this context, the *p*-value quantifies the statistical probability of observing a difference in age acceleration between diseased patients and healthy controls as extreme as, or more extreme than, the observed data under the null hypothesis of equal biological age rates (*H*_0_ : ΔAA1 = 0). An FDR-corrected *p*-value below the standard significance threshold (*p <* 0.05) indicates that the observed acceleration in biological aging is statistically significant and extremely unlikely to be an artifact of random sample variation. The remarkably small *p*-values reported in Table 2 (reaching as low as 10^*−*134^) provide overwhelming empirical evidence that the physical entropy-weighted model measures true pathological age acceleration.

Notably, our physics-informed non-linear clock identified highly significant age acceleration in HIV-infected patients consistently across multiple datasets (GSE53840, GSE53841, GSE100264, GSE107080, GSE117859, GSE117860, GSE140800, GSE185389, GSE185390). It also captured significant age acceleration in Ischemic Heart Disease (IHD) patients (*p* = 5.49 *×* 10^*−*7^ in GSE62867), Rheumatoid Arthritis patients (GSE137593, GSE137594), and Multiple Sclerosis patients (GSE43976).

Furthermore, in blood-cell-count-adjusted age acceleration (AA2) analysis, the Stacy Gradient Boosting Clock demonstrated statistically significant associations with Werner Syndrome (WS) (GSE131752, *p <* 0.05 after FDR correction), a premature aging disorder characterized by genomic instability and accelerated epigenetic drift. It also identified significant AA2 associations for HIV infection across multiple cohorts (GSE67705, GSE67751, GSE77696, GSE143942), confirming its robust performance after adjusting for confounders. The p-value distributions and significant disease associations are shown in Figure 3(a) and (b).

**Figure 3:**
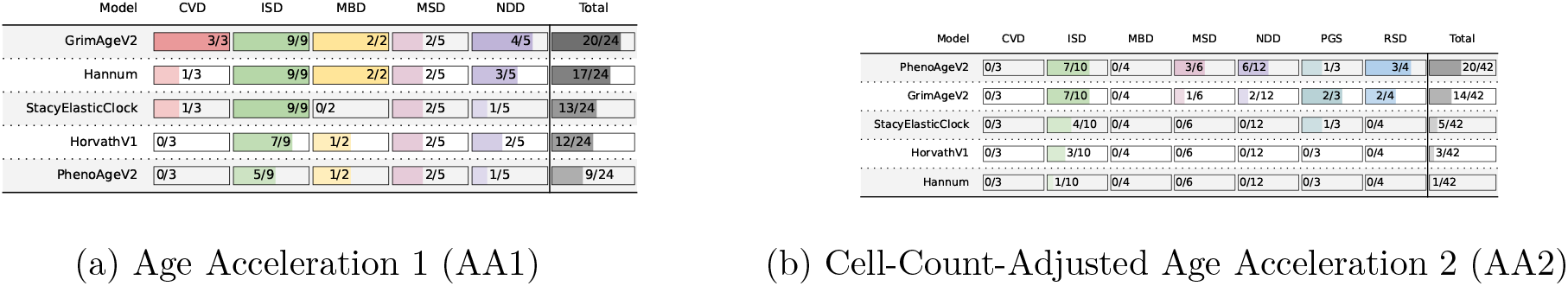
FDR-corrected statistical significance of disease pathology associations with epigenetic Age Acceleration across independent GSE cohorts: (a) raw residual Age Acceleration 1 (AA1), and (b) blood-cell-count-adjusted Age Acceleration 2 (AA2).

## 5 Conclusion

In this study, we introduced the Stacy Gradient Boosting Clock, a physics-informed epigenetic clock that bridges statistical thermodynamics with non-linear machine learning. Traditional epigenetic clocks rely almost exclusively on empirical regression directly over DNA methylation *β*-values, treating cellular aging as a black-box machine learning problem. By contrast, our framework grounds methylome degradation in physical principles by modeling cell-to-cell methylation variability across CpG sites using a reparameterized three-parameter Generalized Gamma Distribution (GGD). Through exact analytical derivation of the GGD Shannon entropy *H*^*cpg*^, we systematically transform raw methylation levels into physically weighted entropy features (*X*^*entropy*^ = *x · H*^*cpg*^), converting raw statistical measurements into indicators of local thermodynamic disorder.

Our extensive benchmarking on the ComputAge repository—comprising 7,419 training samples and 63,001 shared CpG sites across 111 independent studies demonstrates the superior predictive power of this physical formulation. The Stacy Gradient Boosting Clock achieved an outstanding Mean Absolute Error (MAE) of 3.74 years on independent test cohorts. This represents a 30.1% error reduction compared to the gold-standard multi-tissue Horvath clock (MAE = 5.35 years), while dramatically outperforming other established clocks including Hannum (MAE = 7.48 years), PhenoAgeV2 (MAE = 7.60 years), and GrimAgeV2 (MAE = 9.80 years). Furthermore, our clock maintained a low prediction bias with a Median Error of 2.41 years, demonstrating robust generalization across diverse tissues, ages, and profiling platforms.

Crucially, our findings prove that combining Stacy GGD entropy weighting with non-linear gradient boosting (LightGBM) represents a highly effective paradigm for epigenomics. Physical entropy weighting acts as a biophysical prior, highlighting CpG loci experiencing active epigenetic drift while preserving physical scale. Pairing these physical features with decision-tree gradient boosting overcomes a fundamental limitation of traditional ElasticNet clocks: the invalid assumption of linear, non-interacting CpG contributions. LightGBM seamlessly captures complex, high-order non-linear feature interactions and non-monotonic aging trajectories across entropy-weighted loci, unlocking latent predictive signals that linear models miss.

Beyond chronological age accuracy, we validated the biological interpretability of our model through systematic analysis of epigenetic Age Acceleration (AA1 and AA2). The age acceleration derived from physical entropy features exhibited powerful, statistically significant associations with major pathologies after stringent False Discovery Rate (FDR) correction. Notably, our clock identified consistent, highly significant age acceleration in HIV infection across nine independent datasets (achieving *p*-values as strong as 10^*−*134^), as well as Ischemic Heart Disease (*p* = 5.5 *×* 10^*−*7^), Rheumatoid Arthritis, and Multiple Sclerosis. In cell-count-adjusted AA2 analyses, the clock successfully identified accelerated biological aging in Werner Syndrome, a monogenic progeroid disorder characterized by genomic instability and accelerated epigenetic decay.

In summary, this work establishes that incorporating statistical physics and thermodynamic entropy into non-linear gradient boosting models is a superior approach for epigenetic clock construction. By transforming empirical measurements into biophysically meaningful entropy features, the Stacy Gradient Boosting Clock enhances both prediction accuracy and clinical interpretability, offering a powerful, physically grounded biomarker for aging research, disease risk stratification, and longevity intervention trials.

## Acknowledgements

This research was funded in part by the Slovak Research and Development Agency under contracts No. APVV-22-0508, No. APVV-18-0526, by the Slovak Grant Agency for Science under grant VEGA 1/0674/23, and the Cultural and Educational Grant Agency under grant KEGA 006TUKE-4/2024.

## Appendix A: Methylation Entropy Calculation

In this appendix, we provide the detailed step-by-step mathematical derivation of the thermodynamic and Shannon entropy for the reparameterized Generalized Gamma Distribution (GGD), deriving Eq. (8) and Eq. (10).

Starting from the definition of Shannon entropy and thermodynamic entropy:

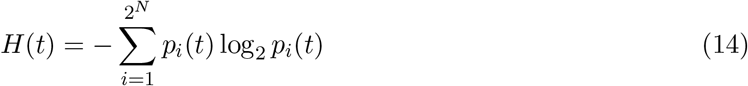

and the reparameterized GGD probability density function:

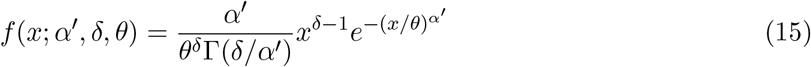

the continuous thermodynamic entropy deviation Δ*S* = *S − S*_0_ (taking reference entropy *S*_0_ = 0) is defined by the continuous integral:

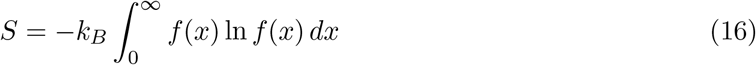

Taking the natural logarithm of the density *f* (*x*; *α*^*′*^, *δ, θ*):

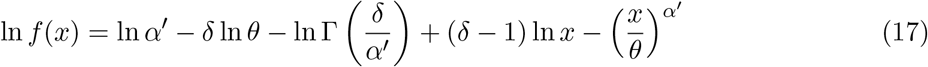

Substituting (17) into the entropy integral formula (16):

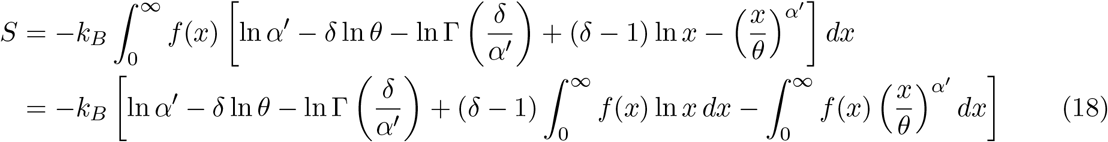

Using the expected values and logarithmic moments of the Generalized Gamma Distribution:

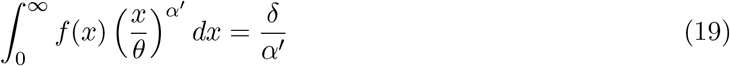

and

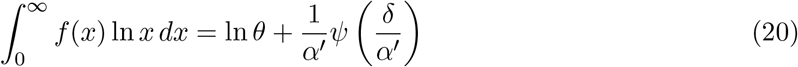

where 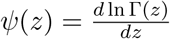 is the digamma function.

Substituting these expectations back into (18):

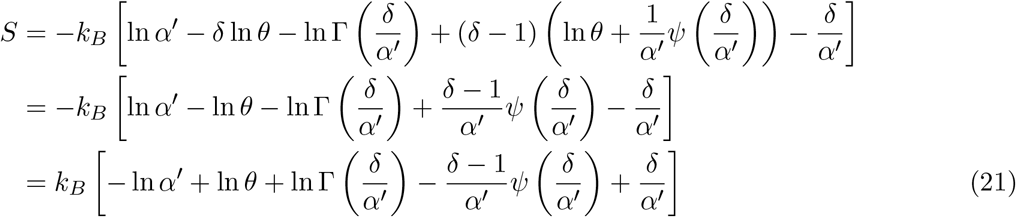

Grouping the logarithmic scale and gamma function terms:

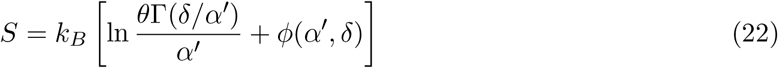

where the auxiliary function *ϕ*(*α*^*′*^, *δ*) is defined as:

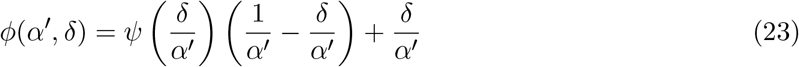

Finally, converting thermodynamic entropy *S*(*t*) to Shannon entropy *H*(*t*) (base 2) via 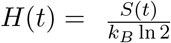 yields:

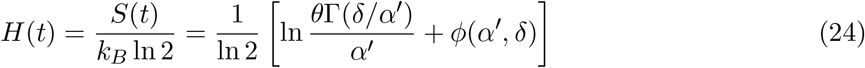

which completes the exact analytical derivation of Eq. (10).

## References

[1] S. Horvath, “DNA methylation age of human tissues and cell types,” Genome Biology, vol. 14, no. 10, p. R115, 2013.

[2] S. Horvath, “Erratum to: DNA methylation age of human tissues and cell types,” Genome Biology, vol. 16, no. 1, p. 96, 2015.

[3] G. Hannum et al., “Genome-wide methylation profiles reveal quantitative views of human aging rates,” Molecular Cell, vol. 49, no. 2, pp. 359–367, 2013.

[4] R. Sánchez and S. A. Mackenzie, “Information thermodynamics of DNA methylome,” PLoS ONE, vol. 11, no. 3, p. e0150427, 2016.

[5] R. Sánchez and S. A. Mackenzie, “DNA methylation dynamics as a thermodynamic process,” Scientific Reports, vol. 13, p. 8914, 2023.

[6] H. Nasrolahpour et al., “Age Formula from DNA Methylation Entropy,” preprint, 2025.

[7] M. Pellegrini et al., “Epigenetic entropy as a biophysical biomarker of aging,” Journal of Theoretical Biology, vol. 579, p. 111697, 2024.

[8] E. W. Stacy, “A generalization of the gamma distribution,” Annals of Mathematical Statistics, vol. 33, no. 3, pp. 1187–1192, 1962.

[9] Computational Aging Lab, “ComputAge: A benchmarking platform for epigenetic clocks,” available at https://github.com/ComputationalAgingLab/ComputAge.

[10] M. E. Levine et al., “An epigenetic biomarker of aging for lifespan and healthspan,” Aging (Albany NY), vol. 10, no. 4, pp. 573–591, 2018.

[11] A. T. Lu et al., “DNA methylation GrimAge strongly predicts lifespan and healthspan,” Aging (Albany NY), vol. 11, no. 2, pp. 303–327, 2019.

[12] G. Ke et al., “LightGBM: A highly efficient gradient boosting decision tree,” in Advances in Neural Information Processing Systems (NeurIPS), vol. 30, pp. 3146–3154, 2017.

[13] W. Zhou et al., “SeSAMe: reducing artifactual detection of DNA methylation by high-density beads in Infinium arrays,” Nucleic Acids Research, vol. 46, no. 20, p. e123, 2018.

[14] M. Raissi et al., “Physics-informed neural networks: A deep learning framework for solving forward and inverse problems involving nonlinear partial differential equations,” Journal of Computational Physics, vol. 378, pp. 686–707, 2019.

[15] R. H. Byrd et al., “A limited memory algorithm for bound constrained optimization,” SIAM Journal on Scientific Computing, vol. 16, no. 5, pp. 1190–1208, 1995.

[16] Y. Benjamini and Y. Hochberg, “Controlling the false discovery rate: a practical and powerful approach to multiple testing,” Journal of the Royal Statistical Society: Series B (Methodological), vol. 57, no. 1, pp. 289–300, 1995.

